# A comprehensive LFQ benchmark dataset on modern day acquisition strategies in proteomics

**DOI:** 10.1101/2021.11.24.469852

**Authors:** Bart Van Puyvelde, Simon Daled, Sander Willems, Ralf Gabriels, Anne Gonzalez de Peredo, Karima Chaoui, Emmanuelle Mouton-Barbosa, David Bouyssié, Kurt Boonen, Christopher J. Hughes, Lee A. Gethings, Yasset Perez-Riverol, Nic Bloomfield, Stephen Tate, Odile Schiltz, Lennart Martens, Dieter Deforce, Maarten Dhaenens

## Abstract

In the last decade, a revolution in liquid chromatography-mass spectrometry (LC-MS) based proteomics was unfolded with the introduction of dozens of novel instruments that incorporate additional data dimensions through innovative acquisition methodologies, in turn inspiring specialized data analysis pipelines. Simultaneously, a growing number of proteomics datasets have been made publicly available through data repositories such as ProteomeXchange, Zenodo and Skyline Panorama. However, developing algorithms to mine this data and assessing the performance on different platforms is currently hampered by the lack of a single benchmark experimental design. Therefore, we acquired a hybrid proteome mixture on different instrument platforms and in all currently available families of data acquisition. Here, we present a comprehensive Data-Dependent and Data-Independent Acquisition (DDA/DIA) dataset acquired using several of the most commonly used current day instrumental platforms. The dataset consists of over 700 LC-MS runs, including adequate replicates allowing robust statistics and covering over nearly 10 different data formats, including scanning quadrupole and ion mobility enabled acquisitions. Datasets are available via ProteomeXchange (PXD028735).

## Background & Summary

Hypothesis-driven biochemical assays have been the foundation of molecular biology for well over a century, with great success. However, the lack of a more holistic view on the biomolecular complexity requires trial-and-error experimentation. Therefore, the past few decades were characterized by a shift towards an experimental design wherein a broader biomolecular perspective of the system is first generated in order to contextualize the hypothesis and the targeted biochemical assays beforehand. These “omics” approaches were enabled by two technical revolutions, i.e. the sequencing of nucleotides and the accurate mass measurement of biomolecules by mass spectrometry (MS).

In its barest form, the output from an MS instrument is merely a list of m/z’s with intensities measured at very precise moments in time. However, MS is quickly evolving towards capturing the full complexity of a biological sample. To this end, not only the accuracy of instruments has improved greatly, they now also incorporate analytical techniques that select or separate analytes based on other physico-chemical properties. In proteomics nowadays, a mass spectrometer thus rarely only measures the *m/z* coordinate and intensity of (fragment) ions. The ion coordinates are mostly supplemented with precursor *m/z*, retention time (*t_R_*) and/or ion mobility coordinates, depending on acquisition strategy. This creates a multidimensional data matrix that captures the complexity of the sample to an unprecedented depth ^1^.

The field of mass spectrometry has diversified greatly, driven by a fast sequence of innovations from many vendors. Modern MS instruments allow the manipulation of ions in countless of different ways, including different ionization methodologies, fragmentation techniques, multipoles, time-of-flight tubes, ion mobility separation devices and trap designs, including the now very dominant orbitrap. The way in which these different ion manipulations are combined has ballooned the number of different acquisition strategies available to the end user today but all these instrumental and strategic innovations are futile if no data analysis pipeline is available to translate the data back into biology. For bottom-up proteomics this implies reconstructing the peptide backbone sequences from their fragment ions because the latter encompasses the specificity for identifying the hundreds of millions of different protein sequences that make up the biotic world.

Conventionally, MS instruments have been operated using data dependent acquisition (DDA) wherein the data from a precursor scan at low energy is used to pinpoint potentially interesting analytes which are then sequentially selected for fragmentation at high energy. These fragmentation spectra can then be identified by a plethora of different algorithms ^2,3^. Data-independent acquisition (DIA) however, is the more intuitive way of analyzing a sample, because it captures all (fragment) ions without any instrumental bias. However, interpreting such complex data matrices has proven difficult and an additional separation dimension, such as ion mobility, was added to increase the discriminating power ^4–8^. Alternatively, configuring a quadrupole to sequentially scan the entire mass range - while still operating “data independent” - alleviates the complexity of the resulting fragmentation spectra even more ^9,10^. This has opened up the way for the many different spectrum-centric and peptide-centric data analysis strategies available today ^11–16^. The latest reduction in complexity or “chimericy” of DIA spectra encompasses continuously scanning the quadrupole as is done with SONAR ^17^ and Scanning SWATH ^18^ and combining quadrupole selection and ion mobility separation, as is done with diaPASEF ^19^. Unsurprisingly, machine learning is taking center stage in mining the various resulting data architectures ^20–27^.

Here, we created a comprehensive dataset on a single benchmark experimental design adapted from Navarro et al. ^28^. It contains a ground truth that serves as a quality control for bioinformatics algorithm development and evaluation. This sample was acquired with adequate replicates on many of the current day instrumental platforms – partially in nano flow LC and partially in capillary flow LC - by most of the available acquisition strategy families, covering all commonly measured ion coordinates (**Figure 1**). Far from being complete, it still is the most comprehensive repository of its kind for algorithmic development and validation, both at the level of identification and quantification. Instead of being yet another way of attaining the highest number of identified peptides, we hope it becomes a resource for compatibility assessment and data analysis quality control. Above all, it is a snapshot of current day completeness of our digital image of the protein world.

**Figure 1.**
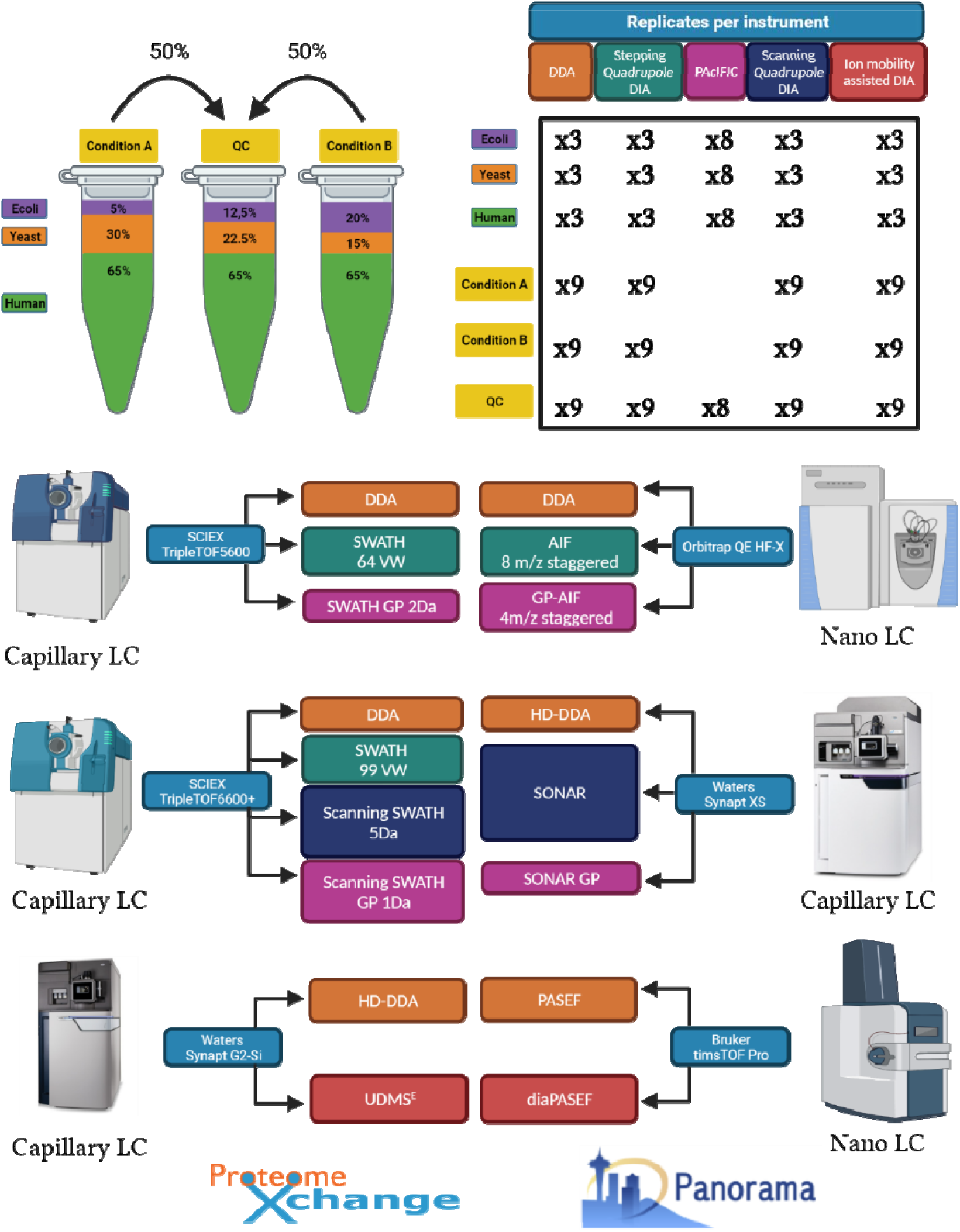
Schematic overview of the different acquisition strategies/instruments applied in the study. A comprehensive LC-MS/MS dataset was generated using samples composed of commercial Human, Yeast and E.coli full proteome digests. Two hybrid proteome samples A and B containing known quantities of Human, Yeast and E.coli tryptic peptides, as described by Navarro et al. were prepared in three consecutive times to include handling variability. Additionally, a QC sample was created by mixing one sixth of each of the six master batches (65% w/w Human, 22.5% w/w Yeast and 12.5% w/w E.coli). These commercial lysates were measured individually and as triple hybrid proteome mixtures each in triplicate using DDA and DIA acquisition methodologies available on six LC-MS/MS platforms, i.e. SCIEX TripleTOF5600 and TripleTOF 6600+, Thermo Orbitrap QE HF-X, Waters Synapt G2-Si and Synapt XS and Bruker timsTOF Pro. The complete dataset was made publicly available to the proteomics community through ProteomeXchange with dataset identifier: PXD028735. In addition, a system suitability workflow (AutoQC) was incorporated on each instrument using commercial E.coli lysate digest which were acquired at multiple timepoints throughout each sample batch. The AutoQC data was automatically imported in Skyline and uploaded to the Panorama AutoQC server using AutoQC loader, enabling system suitability assessment of each LC-MS/MS system used in the dataset.

## Methods

### Sample preparation

Mass spectrometry-compatible Human (P/N: V6951) and Yeast (P/N: V7461) protein digest extracts were purchased from Promega (Madison, Wisconsin, United States). Lyophilised MassPrep E.coli digest standard (P/N:186003196) was purchased from Waters Corporation (Milford, Massachusetts, United States). The extracts were reduced with dithiothreitol (DTT), alkylated with iodoacetamide (IAA) and digested with sequencing grade Trypsin(-Lys C) by the respective manufacturers. The digested protein extracts were reconstituted in a mixture of 0.1% Formic acid (FA) in water (Biosolve B.V, Valkenswaard, The Netherlands) and spiked with iRT peptides (Biognosys, Schlieren, Switzerland) at a ratio of 1:20 v/v. Two master samples A and B were created similar to Navarro et al., each in triplicate, as shown in **Figure 1**. Sample A was prepared by mixing Human, Yeast and E.coli at 65%, 30% and 5% weight for weight (w/w), respectively. Sample B was prepared by mixing Human, Yeast and E.coli protein digests at 65%, 15%, 20% w/w, respectively. The resulting samples have logarithmic fold changes (log2FCs) of 0,-1 and 2 for respectively Human, Yeast and E.coli. One sixth of each of the triplicate master batches of A and B were mixed to create a QC sample, containing 65% w/w Human, 22.5% w/w Yeast and 12.5% w/w E.coli.

### LC-MS/MS

In this section, a detailed description of the different LC-MS/MS parameters is given for each LC-MS/MS instrumental setup applied to generate this comprehensive dataset. All instruments were operated according to the lab’s best practice, i.e. not necessarily the best attainable, but rather most realistic data quality. Sample load was chosen based on LC setup (nano flow = 1 μg on column vs capillary flow = 5 μg on column) and instrument sensitivity. Thus, differences in absolute number of identified peptides and proteins can be attributed to sample load, LC flow rate, MS instrumentation, operator’s choices and search algorithmic compatibility; direct conclusions on MS instrument performance can therefore not be drawn from this dataset.

As a rule of thumb, data-dependent acquisition (DDA) methods use high energy fragmentation spectra (MS2) of narrow mass selections for identification and use the area under the curve of the precursor (MS1) for quantification. Therefore, a cycle time needs to be attained wherein enough datapoints across the precursor elution peak are sampled for accurate quantification. In most data-independent acquisition (DIA) strategies, a broader precursor selection window is used and both identification and quantification can be done at the fragment level, taken that the cycle time for both MS1 and MS2 is adapted to the LC gradient. Finally, Precursor Acquisition Independent From Ion Count (PAcIFIC) is a method that is acquired solely to extend the size of the peptide library for detecting peptides in DIA data and is therefore not strictly dependent on the cycle time. Of note, by scanning the quadrupole instead of acquiring different mass windows separately, acquisition strategies like SONAR and Scanning SWATH create an additional dimension in the data matrix, akin to how ion mobility separation is perceived. Since these are similar to PAcIFIC acquisition, we also acquired gas phase (GP) fractions in SONAR and Scanning SWATH for library building, i.e. with no emphasis on cycle time.

#### 1) SCIEX TripleTOF 5600 (Capilary flow LC)

A TripleTOF 5600 mass spectrometer (Sciex, Concord, Ontario, Canada) fitted with a Duospray ion source operating in positive ion mode, was coupled to an Eksigent NanoLC 400 HPLC system (Eksigent, Dublin, CA). 5 μL of each sample was loaded at 5μL/min with 0.1% Trifluoroacetic acid (TFA) in water and trapped on a YMC TriArt C18 guard column (id 500μm, length 5mm, particle size 3 μm) for 5 minutes. Peptides were separated on a microLC YMC TriArt C18 column (id 300 μm, length 15 cm, particle size 3 μm) maintained at 55°C at a flow rate of 5μL/min by means of trap-elute injection. Mobile phase A consisted of UPLC-grade water with 0.1% (v/v) FA and 3% (v/v) DMSO, and mobile phase B consisted of UPLC-grade ACN with 0.1% (v/v) FA. Peptide elution was performed at 5μL/min using the following gradient: i) 2% to 30% mobile phase B in 120 min, ii) ramp to 90% mobile phase B in 1 min. The washing step at 90% mobile phase B lasted 4 min and was followed by an equilibration step at 2% mobile phase B (starting conditions) for 10 min. Ion source parameters were set to 5.5 kV for the ion spray voltage, 30 psi for the curtain gas, 13 psi for the nebulizer gas and 80°C as source temperature.

##### a. Data-Dependent Acquisition

For DDA (a cycle time of 3.5 s), MS1 spectra were collected between 399-1200 m/z for 500 ms. The 20 most intense precursors ions with charge states 2-5 that exceeded 250 counts per second were selected for fragmentation, and the corresponding fragmentation MS2 spectra were collected between 50-2000 m/z for 151 ms. After the fragmentation event, the precursor ions were dynamically excluded from reselection for 20 s.

##### b. PAcIFIC

For PAcIFIC (a cycle time of 4 s), the TripleTOF5600 was configured to acquire eight gas phase fractionated acquisitions with isolation windows of 4 m/z using an overlapping window pattern from narrow mass ranges, as described by Searle et al (i.e., 396.43 – 502.48; 496.48 – 602.52;596.52 – 702.57;696.57 – 802.61; 796.61 – 902.66; 896.6 – 1002.70; 996.70 – 1102.75; 1096.75 – 1202.80) ^29^. See **Supplementary Table 2** for the actual windowing scheme. MS2 spectra were collected in high-sensitivity mode from 360-1460 m/z, for 75 ms. An MS1 survey scan was recorded per cycle from 360-1460 m/z for 50ms.

##### c. SWATH 64 variable windows

For SWATH (a cycle time of 3.4 s), a 64 variable window acquisition scheme as described by Navarro et al. was used for all samples (**Supplementary Table 1**) ^28^. Briefly, SWATH MS2 spectra were collected in high-sensitivity mode from 50-2000 m/z, for 50 ms. Before each SWATH MS cycle an additional MS1 survey scan in high sensitivity mode from 400-1200 m/z was recorded for 150 ms.

#### 2) SCIEX TripleTOF 6600+ (Capilary flow LC)

A TripleTOF 6600+ mass spectrometer (Sciex, Concord, Ontario, Canada) fitted with an Optiflow ion source operating in positive ion mode, was coupled to an Eksigent NanoLC 425 HPLC system (Eksigent, Dublin, CA). 5 μL of each sample was loaded at 5μL/min with 0.1% FA in water by means of direct injection. Peptides were separated on a Phenomenex Luna Omega Polar C18 column (150 x 0.3 mm, particle size 3 μm) at a column temperature of 30°C. Mobile phase A consisted of UPLC-grade water with 0.1% (v/v) FA, and mobile phase B consisted of UPLC-grade ACN with 0.1% (v/v) FA. Peptide elution was performed at 5μL/min using the following gradient: i) 2% to 30% mobile phase B in 120 min, ii) ramp to 90% mobile phase B in 1 min. The washing step at 90% mobile phase B lasted 4 min and was followed by an equilibration step at 2% mobile phase B (starting conditions) for 10 min. Ion source parameters were set to 4.5 kV for the ion spray voltage, 25 psi for the curtain gas, 10 psi for nebulizer gas (ion source gas 1), 20 psi for heater gas (ion source gas 2) and 100°C as source temperature.

##### a. Data-Dependent Acquisition

For DDA acquisition (a cycle time of 3.3 s), MS1 spectra were collected between 400-1200 m/z for 250 ms. The 30 most intense precursor ions with charge states 2-4 that exceeded 300 counts per second were selected for fragmentation, and the corresponding fragmentation MS2 spectra were collected between 100-1500 m/z for 100 ms. After the fragmentation event, the precursor ions were dynamically excluded from reselection for 10 s.

##### b. SWATH 99 Variable windows

For SWATH (a cycle time of 4 s), a 99 variable window acquisition scheme was used (see **Supplementary Table 3**) ^30^. Briefly, SWATH MS2 spectra were collected in high sensitivity mode from 100-1500 m/z, for 37.5 ms. Before each SWATH MS cycle an additional MS1 survey scan in high sensitivity mode was recorded for 250 ms.

##### c. Scanning SWATH GP 1Da

A Scanning SWATH beta version was installed on the Analyst TF control software in collaboration with Sciex Research. Scanning SWATH Q1 calibration was confirmed by directly infusing a tuning solution (ESI Positive Calibration Solution for the SCIEX X500B System - P/N: 5049910) and by acquiring a pre-built calibration batch (SSCalibration.dab). Afterwards, the calibration was verified by i) running a verification calibration batch and inspect the data in PeakView. Calibration of the Q1 after initial calibration was controlled automatically by internal recalibration of each data file at run time removing the need for subsequent calibration events.

The gas-phase fractionation approach usually acquired in PAcIFIC, were also acquired by Scanning SWATH because this uniquely allows to apply DIA annotation algorithms for library building of subsequent full mass range DIA acquisition. Precursor isolation window was set to 1 m/z and a mass range of 100 m/z was covered in 6 s (average accumulation time per precursor: 59.57 ms). An MS1 scan was included and data was acquired in high resolution mode. Raw data was converted to standard SCIEX data files with an effective precursor isolation of 0.2 m/z bins and Q1 calibration was obtained by running rawSSProcessor.exe.

##### d. Scanning SWATH Fixed 5Da Window

The precursor mass range of 400-900 m/z was covered in 4 s with a 5 m/z isolation window (average accumulation time per precursor: 37.5 ms). A TOF MS scan was included and data was acquired in high sensitivity mode. Raw data was converted to standard SCIEX data files with an effective precursor isolation of 1 m/z bins and Q1 calibration was obtained by running the rawSSProcessor.exe.

#### 3) Thermo Orbitrap QE HF-X (Nano flow LC)

A Thermo Orbitrap QE HF-X (Thermo Fisher Scientific, Waltham, Massachusetts, United States) was coupled to an UltiMate 3000 LC-system (NCS-3500RS Nano/Cap System, Thermo Fisher Scientific). Peptides were separated on an Acclaim PepMap C18 column (id 75 μm, length 50 cm, particle size 2 μm, Thermo Fisher Scientific ref 164942) at a flow rate of 350 nL/min by means of trap-elute injection (Acclaim PepMap C18, id. 300 μm x 5mm) after 3min desalting on a nano-trap cartridge (id.300 μm, length 5mm, Thermo Fisher Scientific ref 160454).

Mobile phase A consisted of UPLC-grade water with 0.1% (v/v) FA, and mobile phase B consisted of UPLC-grade ACN with 0.1% (v/v) FA. Peptide elution was performed at 350 nL/min using the following gradient: i) 2% to 30% mobile phase B in 120 min, ii) ramp to 90% mobile phase B in 1 min. The washing step at 90% mobile phase B lasted 4 min and was followed by an equilibration step at 2% mobile phase B (starting conditions) for 21 min.

##### a. Data-Dependent Acquisition

The data-dependent acquisition runs on the Q Exactive HF-X were acquired with MS survey scans (350-1400 m/z) at a resolution of 60,000, and an AGC target of 3e6. The 12 most intense precursor ions, were selected for fragmentation by high-energy collision–induced dissociation, and the resulting fragments were analyzed at a resolution of 15,000 using an AGC target of 1e5 and a maximum fill time of 22 ms. Dynamic exclusion was used within 30 s to prevent repetitive selection of the same peptide.

##### b. Narrow Window Gas-Phase fractionation (GP) DIA

Narrow-window GP-DIA data was acquired as described by Searle et al ^29^. Briefly, 6 GP runs (400-500,…, 900-1000 m/z) using staggered 4□m/z DIA spectra (4□m/z precursor isolation windows at 30,000 resolution, AGC target 1e6, maximum inject time 60□ms, NCE 27, +3H assumed charge state) were acquired using an overlapping window pattern, described by Pino et al ^31^. In each run, full MS scans matching each part of the fractionated mass range (i.e., either 395-505, 495-605, 595-705, 695-805, 795-905, or 895-1005 m/z), acquired at a resolution of 60,000 using an AGC target of 1e6 and a maximum inject time of 60ms, were interspersed every 25 MS/MS spectra.

##### c. All Ion Fragmentation (AIF)

The AIF DIA data was acquired using a staggered pattern of 75×8 m/z isolation windows over the mass range 400-1000 m/z as described by Pino et al (**Supplementary Table 4**) ^31^. DIA MS/MS scans were acquired at 15,000 resolution, with an AGC target of 1e6, a maximum inject time 20ms, and a NCE of 27. Full MS scans over the range 390-1010 m/z at 60,000 resolution, AGC target 1e6, maximum inject time 60 ms were interspersed every 75 MS/MS spectra.

#### 4) Waters Synapt G2-Si (Capillary flow LC)

An M-class LC system (Waters Corporation, Milford, MA) was equipped with a 1.7 μm CSH 130 C18 300 μm x 100 mm column, operating at 5 μL/min with a column temperature of 55 °C. Mobile phase A was UPLC-grade water containing 0.1% (v/v) FA and 3% DMSO, mobile phase B was ACN containing 0.1% (v/v) FA. Peptides were separated using a linear gradient of 3-30% mobile phase B over 120 minutes. All experiments were conducted on a Synapt G2-Si mass spectrometer (Waters Corporation, Wilmslow, UK). The ESI Low Flow probe capillary voltage was 3 kV, sampling cone 60 V, source offset 60 V, source temperature 80 °C, desolvation temperature 350 °C, cone gas 80 L/hr, desolvation gas 350 L/hr, and nebulizer pressure 2.5 bar. A lock mass reference signal of GluFibrinopeptide B (m/z 785.8426) was sampled every 30 s.

##### a. HD-DDA

Data was acquired according to Helm et al. with minor adaptations ^7^. Briefly, in data-dependent mode, the MS automatically switches between MS survey and MS/MS scans based upon a set of switching criteria, including ion intensity and charge state. Full scan MS and MS/MS spectra (m/z 50 - 5000) were acquired in sensitivity mode. MS survey spectra were acquired using a fixed acquisition time of 250 ms and the ions present in each scan were monitored for the following criteria: more than 3000 intensity/sec and only 2,3,4,5+ charge states. Once criteria were satisfied, the precursor ion isolation width of the quadrupole was set to 1.0 Th around each precursor sequentially. Tandem mass spectra of up to 12 precursors were generated in the trapping region of the ion mobility cell by using a collisional energy ramp from 6/9 V (low mass 50 Da, start/end) to up to 147/183 V (high mass 5000 Da, start/end), with actual values applied dependent upon the precursor m/z. The MS2 scan time was set to 100 ms and the “TIC stop” parameter was set to 100,000 intensity/s allowing a maximum accumulation time of 300 ms (i.e. up to three tandem MS spectra of the same precursor). IMS wave velocity was ramped from 2400 m/s to 450 m/s (start to end) and the pusher/ion mobility synchronized for singly charged fragment ions in MS/MS spectra, with up to 85% duty cycle efficiency.

##### b. UDMS^e^

Data was acquired according to Distler et al. with minor adaptations ^32^. Briefly, Two data functions were acquired over a mass range of m/z 50 to 2000 in alternating mode, differing only in the collision energy applied to the gas cell. In low-energy MS1 mode, data was collected at a constant gas cell collision energy of 4 eV. In elevated energy MS2 mode, the gas cell collision energy was ramped from 10 to 60 eV according to a collision energy look up table in function of drift time. The spectral acquisition time in each mode was 0.6 s with a 0.015 s interscan delay.

#### 5) Waters Synapt XS (Capilary flow LC)

An M-class LC system (Waters Corporation, Milford, MA) equipped with a 1.7 μm CSH 130 C18 300 μm x 100 mm column, operating at 7 μL/min with a column temperature of 55 °C was coupled to a Synapt XS quadrupole oa-ToF mass spectrometer (Waters Corporation, Wilmslow, UK) operating at a mass resolution of 30000, FWHM. The ESI Low Flow probe capillary voltage was 1.8 kV, sampling cone 30 V, source offset 4 V, source temperature 100 °C, desolvation temperature 300 °C, cone gas disabled, desolvation gas 600 L/hr, and nebulizer pressure 3.5 bar was used. The time-of-flight (TOF) mass analyzer of the mass spectrometer was externally calibrated with a NaCsI mixture from m/z 50 to 1990. A lock mass reference signal of GluFibrinopeptide B (m/z 785.8426) was sampled every two minutes. Mobile phase A was water containing 0.1% (v/v) FA, while mobile phase B was ACN containing 0.1% (v/v) FA. The peptides were eluted and separated with a gradient of 5–40% mobile phase B over 120 minutes.

##### a. HD-DDA

In data-dependent mode, the MS instrument automatically switches between MS survey and MS/MS scans based upon a set of switching criteria, including ion intensity and charge state. Full scan MS and MS/MS spectra (m/z 50 - 5000) were acquired in resolution mode. MS survey spectra were acquired using a fixed acquisition time of 200 ms and the ions present in each scan were monitored for criteria intensity more than 5000 intensity/sec and 2,3,4+ charge states. Once criteria were satisfied, the precursor ion isolation width of the quadrupole was set to 1.0 Th around each precursor sequentially. Tandem mass spectra of up to 15 precursors were generated in the trapping region of the ion mobility cell by using a collisional energy ramp from 6/9 V (low mass 50 Da, start/end) to up to 147/183 V (high mass 5000 Da, start/end), with actual values applied dependent upon the precursor m/z. The MS2 scan time was set to 70 ms and the “TIC stop” parameter was set to 100,000 intensity/s allowing a maximum accumulation time of 100 ms (i.e. up to two tandem MS spectra of the same precursor). IMS wave velocity was ramped from 2450 m/s to 550 m/s (start to end) and the pusher/ion mobility synchronized for singly charged fragment ions in MS/MS spectra, with up to 85% duty cycle efficiency.

##### b. SONAR GP

As for Scanning SWATH, the GP fractionation approach which is usually acquired in PAcIFIC was also analysed by SONAR purely to extend the peptide library. Therefore, the mass scale from m/z 400 to 1200 was divided into 100 Da sections, thus requiring 8 injections for each sample. The quadrupole was continuously scanned from the start mass to end mass of each section and a transmission window of 4 Da was used. In low-energy MS1 mode, data were collected at constant gas cell collision energy of 6 eV. In elevated energy MS2 mode, the gas cell collision energy was ramped with values calculated from the start and end mass of the 100 Da mass range being scanned by the quadrupole, and are shown in **Supplementary Table 5**. The spectral acquisition time in each mode was 0.5 s with a 0.02 s interscan delay.

##### c. SONAR

The quadrupole was continuously scanned between m/z 400 to 900, with a quadrupole transmission width of ~20 Da. Two data functions are acquired in an alternating mode, differing only in the collision energy applied to the gas cell. In low-energy MS1 mode, data were collected at constant gas cell collision energy of 6 eV. In elevated energy MS2 mode, the gas cell collision energy was ramped from 16 to 36 eV (per unit charge). The spectral acquisition time in each mode was 0.5 s with a 0.02 s interscan delay.

#### 6) Bruker TimsTOF Pro (Nano flow LC)

An Acquity UPLC M-Class System (Waters Corporation) was fitted with a nanoEase^™^ M/Z Symmetry C18 trap column (100Å, 5 μm, 180 μm x 20 mm) and a nanoEase^™^ M/Z HSS C18 T3 Column (100Å, 1.8 μm, 75 μm x 250 mm, both from Waters Corporation). The sample was loaded onto the trap column in 2min at 5μl/min in 94% mobile phase A and 6% mobile phase B. Mobile phase A is UPLC-grade water with 0.1% FA, while mobile phase B is 80% ACN with 0.1% FA. The Acquity UPLC M-Class system was coupled online to a TimsTOF Pro via a CaptiveSpray nano-electrospray ion source (Bruker Daltonics, Bremen, Germany), with an ion transfer capillary temperature at 180°C. Liquid chromatography was performed at 40 °C and with a constant flow of 400 nL/min. Peptides were separated using a linear gradient of 2-30% mobile phase B over 120 minutes. The TimsTOF Pro elution voltages were calibrated linearly to obtain reduced ion mobility coefficients (1/K0) using three selected ions of the Agilent ESI-L Tuning Mix (m/z, 1/K_0_: 622.0289 Th, 0.9848 Vs cm^−2^; 922.0097 Th, 1.1895 Vs cm^−2^; 1222.9906 Th, 1.3820 Vs cm^−2^).

##### a. PASEF

Parallel Accumulation–Serial Fragmentation DDA (PASEF) was used to select precursor ions for fragmentation with 1 TIMS-MS scan and 10 PASEF MS/MS scans, as described by Meier et al. in 2018 ^33^. The TIMS-MS survey scan was acquired between 0.70 - 1.45 V.s/cm^2^ and 100 - 1700 *m/z* with a ramp time of 100 ms. The 10 PASEF scans contained on average 12 MS/MS scans per PASEF scan with a collision energy of 10 eV. Precursors with 1 – 5 charges were selected with the target value set to 20 000 a.u and intensity threshold to 2500 a.u. Precursors were dynamically excluded for 0.4 min.

##### b. diaPASEF

The diaPASEF method was implemented as described by Meier et al. in 2019 ^19^. The DIA parameters that define the windows can be found in **Supplementary Table 6**. The DIA range was set to 400-1200 m/z with 16 diaPASEF scans of 25 m/z isolation windows, including an overlap of 1 Da. Each diaPASEF scan consisted of two steps (measuring two 25 Da intervals), with each step spanning an IMS range of 0.3 V.s/cm^2^. The lower IMS value increased linear from 0.6 to 0.834375 for the diaPASEF scans. The TIMS-MS scan was identical to the PASEF method.

### Data Records

Data record 1. The mass spectrometry DDA and DIA-MS proteomics data including instrument raw files (.wiff, .raw, .d) have been deposited to the ProteomeXchange Consortium via the PRIDE partner repository with the dataset identifier, PXD028735 ^34,35^. For every instrument, a separate Sample and Data Relationship File (SDRF) and an Investigation Description File (IDF) have been uploaded to ProteomeXchange. Both the SDRF and IDF file formats are relatively new in Proteomics and were developed in a collaboration between EuBIC and the Proteomics Standards Initiative (PSI) ^36,37^. These files are used to annotate the sample metadata and link the metadata to the corresponding data file(s) and thus will improve the reproducibility and reanalysis of this comprehensive benchmark dataset.

Data record 2. The AutoQC data analysed in Skyline is available from Panorama Public with the link https://panoramaweb.org/LFQBenchmark.url.

### Technical Validation

We continuously performed system suitability procedures to monitor LC-MS/MS performance in a longitudinal fashion. Therefore, we ran an AutoQC complex lysate, i.e. a commercial E.coli protein digest extract (**Supplementary Table 7**), every 3 to 4 samples over all acquired runs on all instruments. All the AutoQC samples were acquired in DDA on each LC-MS/MS instrument, except for the Synapt XS, Orbitrap QE-HF and the timsTOF Pro, where incidentally DDA and DIA acquisitions were alternated. The same mobile phase A and B composition as for the benchmark samples was used as for the benchmark samples, but the gradient applied was modified to reduce the time required to acquire the complete sample batch: linear 3-40% B in 60 minutes, up to 85% B in 2 minutes, isocratic at 85% B for 7 minutes, down to 3% B in 1 minute and isocratic at 3% B for 10 minutes. Note that the timsTOF Pro AutoQC samples were acquired using the 120min gradient similar to the actual hybrid proteome samples.

System suitability assessment was performed by monitoring peptide-identification free metrics (i.e. retention time, peak area, mass accuracy, etc.) extracted with the vendor neutral Panorama AutoQC framework^38,39^. To isolate a set of peptides that can be used for this, triplicate AutoQC samples acquired on each instrument were peak picked using MSConvert (version 3.0.20070) and the corresponding .MGF files were searched against an E.coli FASTA database using Mascot Daemon (v2.7). The searches were performed with following settings: (i) 20 ppm peptide mass tolerance, (ii) 50 ppm fragment mass tolerance and (iii) two allowed missed cleavages. The peptide and protein identification results were exported as Mascot .DAT file and imported into Skyline Daily (version 21.1.1.160). The five highest ranked proteins were retained in the target list and after importing one of the AutoQC .raw files, we manually verified and removed each precursor with co-eluting peptides and low MS1 signal intensity before a Skyline file was saved as template file. Finally, a configuration file for each setup was created with the AutoQC Loader software (version 21.1.0.158) which leads to the automatic import of every sample, with the pattern “AutoQC” in the .raw file or folder structure, in the Skyline template .The data and skyline reports were published to the PanoramaWeb folder “U of Ghent Pharma Biotech Lab - LFQBenchmark across Instrument Platforms” containing six subfolders for each instrumental platform.

For each instrument, peak area, retention time and mass accuracy were manually checked by plotting these metrics in Levey-Jennings plots. For almost every instrument a few outliers were detected, as can be expected on a dataset of over 600 LCMS runs. Fortunately, most of these can be explained by inspecting the raw data and by personal communication with the technicians acquiring the respective datasets. **Figure 2** illustrates one such case. More specifically, two AutoQC samples display a near-complete loss in peak area in the TripleTOF6600+ DDA dataset. Indeed, these were caused by (a) a wrong vial in the sample tray and (b) an empty vial. When these two samples are removed from the QC plot, a more coherent perspective on the variation in standard deviation is seen in the Levey-Jennings plot. Other instances that we have already found include (i) a significant shift in standard deviation in peak area reported for both the Orbitrap, timsTOF Pro and Synapt XS dataset because AutoQC samples were incidentally acquired in two different acquisition methodologies, i.e. in DDA and DIA; (ii) For the timsTOF Pro, a drift in retention time was seen, indicating LC related technical variation which could have been caused by e.g. too short column equilibration times; (iii) In the Orbitrap QE HF-X AutoQC data one peptide (EEAIIK) was undetectable in all the AutoQC samples acquired in AIF. Manual inspection of the acquisition in Skyline (easily accessible through the Panorama QC pipeline) surfaced that it fell out of the precursor m/z range (351.7053) that was acquired.

**Figure 2.**
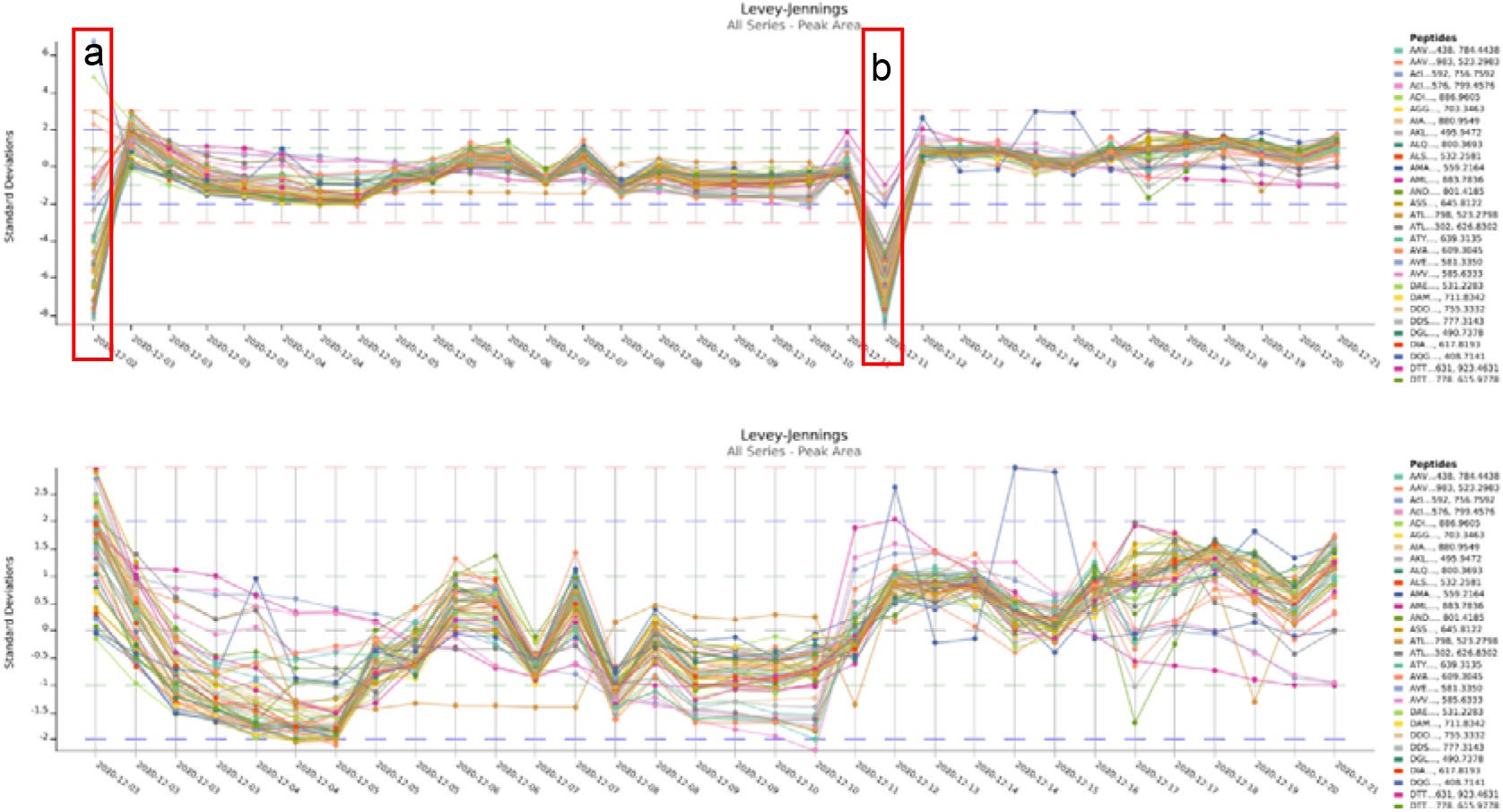
Levey-Jennings plot of the standard deviation in peak area for 50 selected precursors acquired in DDA with the TripleTOF6600+. The upper chart shows two distinct outliers, acquired respectively on the 2^nd^ and 12^th^ of December (red boxes). Manual inspection of the data shows these were caused by **(a)** a wrong vial in the sample tray and **(b)** an empty vial. When these two samples are excluded from the Levey-Jennings plot (lower chart), a significant drop in standard deviation over the time period of data acquisition is seen.

As expected in such a massive MS proteomics experiments, it seems that for every instrument some outliers were recorded, most of which have explanations common to the field. Above all, this demonstrates the necessity of a performant system suitability workflow to increase the reproducibility and quality of LC-MS/MS proteomics datasets^40^.

### Usage Notes

A comprehensive dataset such as the one presented here is inspired by the bioinformatics need to cope with the recent expansion of novel acquisition strategies and data dimensions. Apart from being a repository for validating both performance and compatibility of (new) bioinformatic pipelines, it can serve as a reference for general proteomics courses (e.g. Skyline tutorials, SWATH/DIA course) and be applied for training and validating machine/deep learning algorithms. As such, it is intended to facilitate our understanding of the impact of instrumentation on the perspective that is generated on protein biology and to a larger extent to help unify the field of proteomics.

To demonstrate the applicability of this data repository, we assessed the impact of instrumentation on the most conventional data format acquired by all instruments i.e. DDA. Therefore, for every instrumental platform, triplicate Human, Yeast and E.coli DDA runs were peak picked with MSConvert (version 3.0.21285) and exported as a Mascot .MGF file (**Figure 3A**). MS Convert software was chosen as it is vendor-independent and contains each vendor’s implementation for peak picking, with the exception of UNIFI i.e. Waters. Therefore, Progenesis QI for Proteomics (version 4.2.7207) was used for the Waters DDA data. By doing so, the MS1 precursor space was aligned in the retention time dimension before peak picking. Subsequently, all MS/MS spectra were exported as .MGF file for peptide identification. A standard search with carbamidomethylation of cysteine as fixed modification was performed using a database containing the Human, Yeast and E.coli protein sequences (downloaded from Uniprot on 19/01/2021) and with following parameters: i) mass error tolerances for the precursor ions and the fragment ions were set at 20 ppm and 50 ppm, respectively; ii) enzyme specificity was set to trypsin, allowing up to one missed cleavage. Next, the results were exported as .DAT file and imported into Skyline to create a non-redundant spectral library with BiblioSpec. Afterwards, the .BLIB file was converted to a .dlib and .msp file format respectively using EncyclopeDIA ^29^. The resulting .msp file was converted using a Python conversion tool (speclib_to_mgf.py) built-in MS^2^PIP, to create a peptide record file (.PEPREC) and .MGF file. Next the proportion (amount) of peptide identifications overlapping between the different instruments was assessed using a custom Python script (**Figure 3B**).

**Figure 3.**
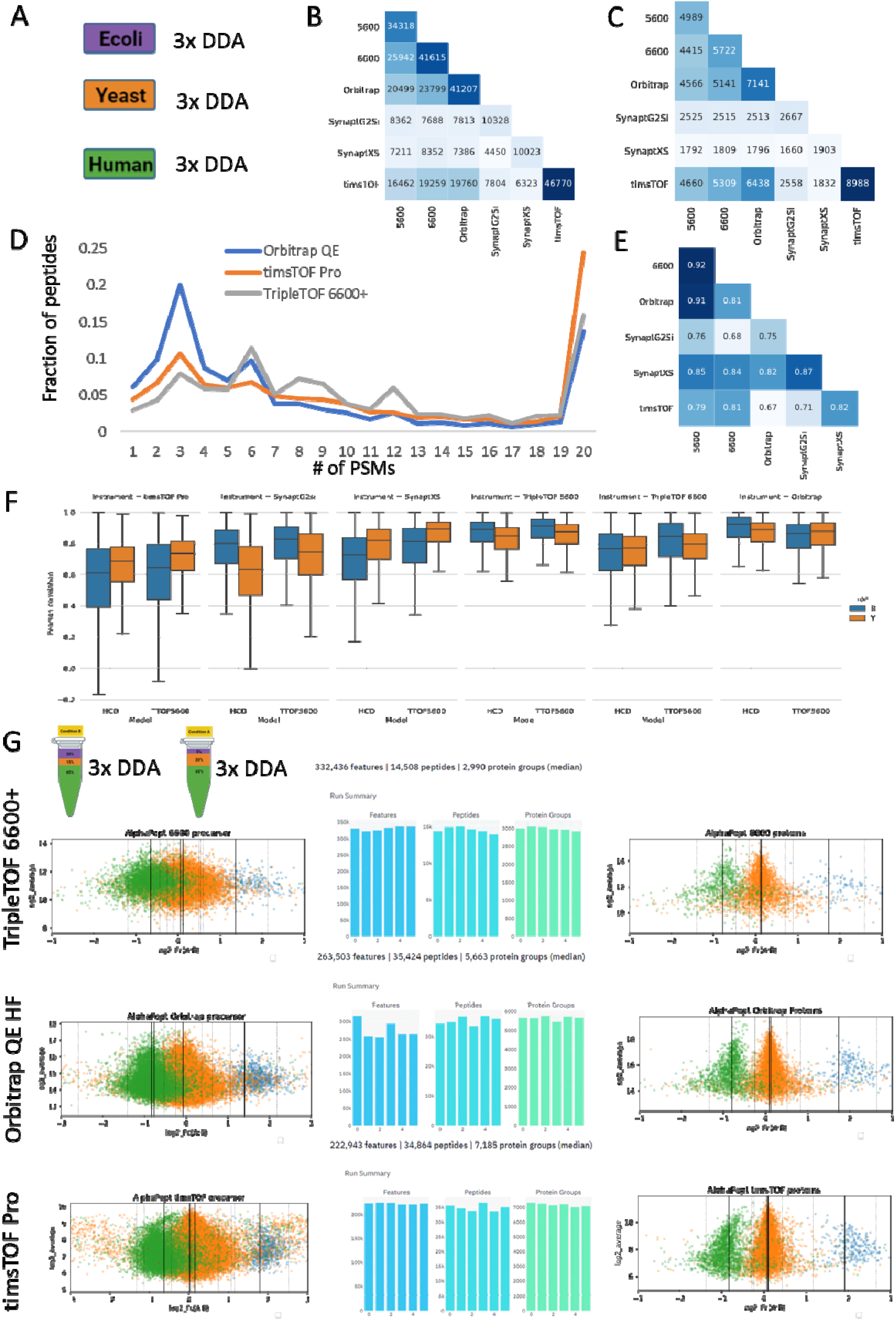
Comparing the DDA data of six different instruments. Experimental design (A) Triplicate measurements of three individual proteomes. (B) The overlap in uniquely identified peptide sequences and (C) proteins between the six instruments. (D) Number of PSMs per peptide identification throughout nine DDA runs on three different proteomes for three instruments. (E) Pearson Correlation Coefficient (PCC) of the fragment intensities were calculated between the shared identified peptides from the DDA replicates between each instrument. The numbers in each box correspond to the median spectrum PCC between the instrument on the x-axis and the instrument on the y-axis. Dark blue color indicates a higher degree of overlap or higher median PCC. (F) Boxplots of the Pearson correlation coefficients (PCC) between the MS^2^PIP predicted (HCD and TTOF5600 model) and experimental fragment intensities across the six different LC-MS instruments. (G) The benchmark design of mixed proteomes for three instruments as annotated and quantified using AlphaPept. Here, triplicate runs of Condition A and Condition B were used, resulting in the six bars depicted in the middle, respectively representing the number of MS1 features, the number of identified peptides and the number of identified proteins for each instrument. The log-fold plots to the left depict the distribution of the peptide ratios in the x-axis as a function of their intensity in the y-axis; protein log fold changes are depicted to the right.

Since each instrument analysed the same commercial protein digests, a large overlap in peptide identifications would be expected. However, while the timsTOF Pro, TripleTOF 6600+ and Orbitrap QE HF-X roughly identify a similar number of peptide sequences (approximately 40,000 over nine LCMS runs), the overlap in identified sequences is in the order of 50%, with that between the Orbitrap QE HF-X and the TripleTOF5600 and 6600+ being overall 10% higher than the overlap with the timsTOF Pro. While at the protein level a lot more coherency is found, it is still striking how the Orbitrap QE and especially the timsTOF Pro have significantly more proteins from a similar number of annotated peptides (**Figure 3C**).

A benchmark like the one presented here however, allows to investigate potential assertions attributed to different stages of the workflow.

#### Acquisition

Even with DDA, the acquisition parameters can be modified substantially. A first, crucial difference is the used flow rate of the LC system: the TripleTOF 6600+ was coupled to a 5μl/min flow, while the Orbitrap and timsTOF Pro used nanoflow (350 and 400 nL/min, respectively). While nano flow is more sensitive, it is also more fragile, which is reflected in some of the quality control samples uploaded to Panorama QC. A second important parameter is dynamic exclusion, i.e. how long a certain precursor is excluded from fragmentation to avoid redundancy. As found in the methods section, this was set to 10s, 24s and 30s for the TripleTOF 6600+, the timsTOF Pro and the Orbitrap QE HF-X, respectively. This parameter usually is a function of LC gradient length and of consistent MSMS spectral quality, which can be facilitated through e.g. automatic gain control (AUC) in orbitrap instruments, but not in QTOF designs. As a result, the orbitrap sampled the peptides most efficiently, with 20% of the peptides only sampled once in a run, i.e. three times over triplicate runs of the same proteome (**Figure 3D**). A third aspect of acquisition is the addition of ion mobility separation (IMS) capabilities in Waters and Bruker instruments. Importantly, they both use IMS in a very different way, forcing a deeper understanding of instrumental architecture and how ion mobility is applied by both vendors in DDA relative to the quadrupole (Q) selection and the collision-induced dissociation (CID). Briefly, the order of ion manipulation is Q-CID-IMS for the so-called High Definition DDA (HD-DDA) in the Waters series and it is IMS-Q-CID in the Bruker instruments. Therefore, Waters separates the fragment ions in IMS and leverages the efficient charge state separation to synchronize the pusher of the TOF tube with singly charged fragment ions in order to only and nearly one hundred percent efficiently sample the single charged fragments ^7^. This can be expected to impact the fragment intensities, especially when detector saturation in MSMS occurs. In the timsTOF Pro on the other hand, the IMS is in fact resolving the precursor ion space before fragmentation, leading to a different selection of the peptide precursor space compared to other devices that do not use IMS (in this way).

#### Raw data

Next, the differences in the fragmentation process between the instruments (all of which are beam-type CID) can be assessed by mutually mapping the MS2 intensities. **Figure 3E** shows their median Pearson correlation coefficients (PCC). By definition, this only plots commonly found peptides. The largest differences in fragment intensities are found between timsTOF Pro and Orbitrap QE HF-X, potentially underlying part of the differences seen in peptide and protein identification. Importantly, the recent introduction of machine learning algorithms has illustrated the potential of fragment intensity prediction for proteomics ^24,25,41–43^. Therefore, to attain an even deeper understanding of these fragmentation differences, we calculated the PCC of both b- and y-ions compared to two prediction models (i.e. HCD and TTOF 5600) from MS^2^PIP ^44^ (**Figure 3F**). This confirms that the orbitrap and triple TOF designs have a very similar fragmentation pattern. Note that the normalized collision energy was not taken into account as input feature ^25,45,46^. Additionally, the Synapt and timsTOF instruments are sampling the ion beam more efficiently, through ion mobility (Waters) or trapping and pusher design (Bruker), potentially causing more frequent detector saturation in the fragment space, leading to skewed intensity patterns compared to the predicted spectra. Indeed, augmented gain controlled in orbitrap devices can efficiently avoid this.

#### Sample Complexity

We also acquired the proteomes as mixtures in known ratios, and selected a triplicate series of Condition A and B for a subsequent analysis (**Figure 3G**). This higher sample complexity is considerably more challenging for DDA acquisition, because the instrument needs to do a faster sampling of the precursor ion space. To create a more comprehensive picture of the resulting data, we turned to AlphaPept, an alternative annotation algorithm that also quantifies the number of features in MS1, i.e. potential peptide species in the sample ^47^. Strikingly, while the TripleTOF 6600+ generated the most features (330k compared to 260k and 220k for the Orbitrap QE HF-X and the timsTOF Pro), it annotated considerably less peptides as well as proteins compared to the two other instruments (**Figure 3G**). Still, the peptide/protein bias persists for the Orbitrap and timsTOF Pro comparison. Importantly, AlphaPept support for Sciex was only preliminarily implemented through mzML preprocessing, potentially underlying the lower annotation rate and making it a perfect example of how the benchmark can support the development of compatibility in the future.

#### Data analysis: Quantification

Next, we used the benchmark design to verify to what extent the annotated peptides and proteins were found in their respective ratios (**Figure 1**). Apart from being a benchmark target for quantification algorithms, this ground truth can also be considered a very rough empirical estimation of the false discovery rate (FDR)^1^. At the peptide level the spread of especially Human peptides into other ratios is increasingly visible in the order TripleTOF 6600+ < Orbitrap QE HF < timsTOF Pro, which alludes to increasing misannotation (**Figure 3G**). While ratio estimation is known to become compromised with decreasing intensities of the MS1 features selected, the relative position of these peptides on the y-axis argues against this intensity explanation. Unfortunately, the exported ion counts of the timsTOF Pro differ at least 1-2 orders of magnitude compared to both the Orbitrap and the TripleTOF 6600+ and more advanced (conversion) algorithms are needed to compare these. However, at the protein level there was no overall skewing in the AlphaPept results that would point towards a significant increase in FDR in any of the instruments.

In conclusion, considerable bioinformatics challenges remain to be tackled before the intricacies of the data from contempory instrumentation and their acquisition parameters can truly be defined.

In conclusion, it is clear from this preliminary data usage case on the simple and most commonly applied DDA strategies that a lot of insights on data structure and bias were still hidden. Especially in light of the more recent DIA strategies, which were not analysed in this Scientific Data report, a lot of work needs to be done before the field truly understands how data acquisition and data analysis effectively perform. Importantly, each step in the data processing can greatly impact the final outcome and we especially anticipate a renewed interest in (multidimensional) peak picking algorithms as a currently underappreciated preprocessing step of the complex DIA data ^1^. Additionally, recent annotation tools have doubled the number of peptides that can be extracted from the same DIA data, an asset not expected for novel DDA annotation algorithms ^21,28^. We would therefore like to invite the developers of all current bioinformatics tools to make use of this comprehensive dataset to benchmark their performance for each instrument individually and adapt their algorithms to increase the performance on all. This dataset was compiled from different labs and therefore also captures differences in instrument usage peculiar to the field, yet it all can be assessed in the Panorama QC metrics that are also publically available. Especially a better understanding of the detectable and annotatable ion space will move the field forward and help researchers make informed decisions on the best acquisition strategies for their application or biological question under investigation.

## Code Availability

MS^2^PIP is open source, licensed under the Apache-2.0 License, and hosted on https://github.com/compomics/ms2pip_c. The Jupyter notebooks used to generate **Figure 3 (B,C,E and F)** are available through Zenodo, under DOI: 10.5281/zenodo.5714380.

## Acknowledgements

This research was funded by grants from the Research Foundation Flanders (FWO) awarded to BVP (grant number: 11B4518N), RG (1S50918N), and MD (12E9716N). Hans Vissers is acknowledged for his assistance with data conversion and formatting. This work was supported in part by the French Ministry of Research with the Investissement d’Avenir Infrastructures Nationales en Biologie et Santé program (ProFi, Proteomics French Infrastructure project; ANR-10-INBS-08).

## Author contributions

BVP, SW, SD and MD conceived the study, BVP performed the TripleTOF 5600 and 6600+ data acquisition, SD performed the Synapt G2-Si data acquisition, AGP, DB, EM and KC performed the Orbitrap data acquisition, KB performed the timsTOF Pro data acquisition. CH and LG performed the Synapt XS data acquisition. BVP and SD prepared the samples and RG and SW wrote the scripts to generate Figure 3. YPR organised the ProteomeXchange submission. NB and ST helped us to set up the acquisition methodologies on the TripleTOF 6600+ and provided valuable insights on the differences seen for each instrument. BVP and MD wrote the draft manuscript with contributions from all authors. MD supervised and DD and LM co-supervised the experiment.

