## Supplementary Information for "A comprehensive LFQ benchmark dataset on modern day acquisition strategies in proteomics"

Table of Contents

Supplementary Table 1. 64 Variable Window Scheme applied for SWATH data acquired on the SCIEX TripleTOF5600

| Start | End | CES |  | Start | End | CES |
| --- | --- | --- | --- | --- | --- | --- |
| 399.5 | 408.2 | 15 |  | 695.9 | 706.9 | 15 |
| 407.2 | 415.8 | 15 |  | 705.9 | 715.9 | 15 |
| 414.8 | 422.7 | 15 |  | 714.9 | 726.2 | 15 |
| 421.7 | 429.7 | 15 |  | 725.2 | 737.4 | 15 |
| 428.7 | 437.3 | 15 |  | 736.4 | 746.6 | 15 |
| 436.3 | 444.8 | 15 |  | 745.6 | 757.5 | 15 |
| 443.8 | 451.7 | 15 |  | 756.5 | 767.9 | 15 |
| 450.7 | 458.7 | 15 |  | 766.9 | 779.5 | 15 |
| 457.7 | 466.7 | 15 |  | 778.5 | 792.9 | 15 |
| 465.7 | 473.4 | 15 |  | 791.9 | 807.0 | 15 |
| 472.4 | 478.3 | 15 |  | 806.0 | 820.0 | 15 |
| 477.3 | 485.4 | 15 |  | 819.0 | 834.2 | 15 |
| 484.4 | 491.2 | 15 |  | 833.2 | 849.4 | 15 |
| 490.2 | 497.7 | 15 |  | 848.4 | 866.0 | 15 |
| 496.7 | 504.3 | 15 |  | 865.0 | 884.4 | 15 |
| 503.3 | 511.2 | 15 |  | 883.4 | 899.9 | 15 |
| 510.2 | 518.2 | 15 |  | 898.9 | 919.0 | 15 |
| 517.2 | 525.3 | 15 |  | 918.0 | 942.1 | 15 |
| 524.3 | 533.3 | 15 |  | 941.1 | 971.6 | 15 |
| 532.3 | 540.3 | 15 |  | 970.6 | 1006.0 | 15 |
| 539.3 | 546.8 | 15 |  | 1005.0 | 1053.0 | 15 |
| 545.8 | 554.5 | 15 |  | 1052.0 | 1110.6 | 15 |
| 553.5 | 561.8 | 15 |  | 1109.6 | 1200.5 | 15 |
| 560.8 | 568.3 | 15 |  |  |  |  |
| 567.3 | 575.7 | 15 |  |  |  |  |
| 574.7 | 582.3 | 15 |  |  |  |  |
| 581.3 | 588.8 | 15 |  |  |  |  |
| 587.8 | 595.8 | 15 |  |  |  |  |
| 594.8 | 601.8 | 15 |  |  |  |  |
| 600.8 | 608.9 | 15 |  |  |  |  |
| 607.9 | 616.9 | 15 |  |  |  |  |
| 615.9 | 624.8 | 15 |  |  |  |  |
| 623.8 | 632.2 | 15 |  |  |  |  |
| 631.2 | 640.8 | 15 |  |  |  |  |
| 639.8 | 647.9 | 15 |  |  |  |  |
| 646.9 | 654.8 | 15 |  |  |  |  |
| 653.8 | 661.5 | 15 |  |  |  |  |
| 660.5 | 670.3 | 15 |  |  |  |  |
| 669.3 | 678.8 | 15 |  |  |  |  |
| 677.8 | 687.8 | 15 |  |  |  |  |
| 686.8 | 696.9 | 15 |  |  |  |  |

Supplementary Table 2. PacIFIC Variable Window Schemes used to acquire the GP-Narrow Window DIA data acquired on the SCIEX TripleTOF5600

| Start | End | CES |  | Start | End | CES |  | Start | End | CES |  | Start | End | CES |
| --- | --- | --- | --- | --- | --- | --- | --- | --- | --- | --- | --- | --- | --- | --- |
| 396.43 | 400.43 | 15 |  | 454.46 | 458.46 | 15 |  | 496.48 | 500.48 | 15 |  | 554.50 | 558.50 | 15 |
| 400.43 | 404.43 | 15 |  | 458.46 | 462.46 | 15 |  | 500.48 | 504.48 | 15 |  | 558.50 | 562.51 | 15 |
| 404.43 | 408.44 | 15 |  | 462.46 | 466.46 | 15 |  | 504.48 | 508.48 | 15 |  | 562.51 | 566.51 | 15 |
| 408.44 | 412.44 | 15 |  | 466.46 | 470.46 | 15 |  | 508.48 | 512.48 | 15 |  | 566.51 | 570.51 | 15 |
| 412.44 | 416.44 | 15 |  | 470.46 | 474.47 | 15 |  | 512.48 | 516.48 | 15 |  | 570.51 | 574.51 | 15 |
| 416.44 | 420.44 | 15 |  | 474.47 | 478.47 | 15 |  | 516.48 | 520.49 | 15 |  | 574.51 | 578.51 | 15 |
| 420.44 | 424.44 | 15 |  | 478.47 | 482.47 | 15 |  | 520.49 | 524.49 | 15 |  | 578.51 | 582.51 | 15 |
| 424.44 | 428.44 | 15 |  | 482.47 | 486.47 | 15 |  | 524.49 | 528.49 | 15 |  | 582.51 | 586.52 | 15 |
| 428.44 | 432.45 | 15 |  | 486.47 | 490.47 | 15 |  | 528.49 | 532.49 | 15 |  | 586.52 | 590.52 | 15 |
| 432.45 | 436.45 | 15 |  | 490.47 | 494.47 | 15 |  | 532.49 | 536.49 | 15 |  | 590.52 | 594.52 | 15 |
| 436.45 | 440.45 | 15 |  | 494.47 | 498.48 | 15 |  | 536.49 | 540.50 | 15 |  | 594.52 | 598.52 | 15 |
| 440.45 | 444.45 | 15 |  | 498.48 | 502.48 | 15 |  | 540.50 | 544.50 | 15 |  | 598.52 | 602.52 | 15 |
| 444.45 | 448.45 | 15 |  |  |  |  |  | 544.50 | 548.50 | 15 |  |  |  |  |
| 448.45 | 452.46 | 15 |  |  |  |  |  | 548.50 | 552.50 | 15 |  |  |  |  |
| 452.46 | 456.46 | 15 |  |  |  |  |  | 552.50 | 556.50 | 15 |  |  |  |  |
| 456.46 | 460.46 | 15 |  |  |  |  |  | 556.50 | 560.50 | 15 |  |  |  |  |
| 460.46 | 464.46 | 15 |  |  |  |  |  | 560.50 | 564.51 | 15 |  |  |  |  |
| 464.46 | 468.46 | 15 |  |  |  |  |  | 564.51 | 568.51 | 15 |  |  |  |  |
| 468.46 | 472.46 | 15 |  |  |  |  |  | 568.51 | 572.51 | 15 |  |  |  |  |
| 472.46 | 476.47 | 15 |  |  |  |  |  | 572.51 | 576.51 | 15 |  |  |  |  |
| 476.47 | 480.47 | 15 |  |  |  |  |  | 576.51 | 580.51 | 15 |  |  |  |  |
| 480.47 | 484.47 | 15 |  |  |  |  |  | 580.51 | 584.52 | 15 |  |  |  |  |
| 484.47 | 488.47 | 15 |  |  |  |  |  | 584.52 | 588.52 | 15 |  |  |  |  |
| 488.47 | 492.47 | 15 |  |  |  |  |  | 588.52 | 592.52 | 15 |  |  |  |  |
| 492.47 | 496.48 | 15 |  |  |  |  |  | 592.52 | 596.52 | 15 |  |  |  |  |
| 496.48 | 500.48 | 15 |  |  |  |  |  | 596.52 | 600.52 | 15 |  |  |  |  |
| 398.43 | 402.43 | 15 |  |  |  |  |  | 498.48 | 502.48 | 15 |  |  |  |  |
| 402.43 | 406.43 | 15 |  |  |  |  |  | 502.48 | 506.48 | 15 |  |  |  |  |
| 406.43 | 410.44 | 15 |  |  |  |  |  | 506.48 | 510.48 | 15 |  |  |  |  |
| 410.44 | 414.44 | 15 |  |  |  |  |  | 510.48 | 514.48 | 15 |  |  |  |  |
| 414.44 | 418.44 | 15 |  |  |  |  |  | 514.48 | 518.49 | 15 |  |  |  |  |
| 418.44 | 422.44 | 15 |  |  |  |  |  | 518.49 | 522.49 | 15 |  |  |  |  |
| 422.44 | 426.44 | 15 |  |  |  |  |  | 522.49 | 526.49 | 15 |  |  |  |  |
| 426.44 | 430.45 | 15 |  |  |  |  |  | 526.49 | 530.49 | 15 |  |  |  |  |
| 430.45 | 434.45 | 15 |  |  |  |  |  | 530.49 | 534.49 | 15 |  |  |  |  |
| 434.45 | 438.45 | 15 |  |  |  |  |  | 534.49 | 538.49 | 15 |  |  |  |  |
| 438.45 | 442.45 | 15 |  |  |  |  |  | 538.49 | 542.50 | 15 |  |  |  |  |
| 442.45 | 446.45 | 15 |  |  |  |  |  | 542.50 | 546.50 | 15 |  |  |  |  |
| 446.45 | 450.45 | 15 |  |  |  |  |  | 546.50 | 550.50 | 15 |  |  |  |  |
| 450.45 | 454.46 | 15 |  |  |  |  |  | 550.50 | 554.50 | 15 |  |  |  |  |

| Start | End | CES |  | Start | End | CES |  |  | Start | End | CES |  | Start | End | CES |
| --- | --- | --- | --- | --- | --- | --- | --- | --- | --- | --- | --- | --- | --- | --- | --- |
| 596.52 | 600.52 | 15 |  | 654.55 | 658.55 | 15 |  |  | 696.57 | 700.57 | 15 |  | 754.59 | 758.59 | 15 |
| 600.52 | 604.52 | 15 |  | 658.55 | 662.55 | 15 |  |  | 700.57 | 704.57 | 15 |  | 758.59 | 762.60 | 15 |
| 604.52 | 608.53 | 15 |  | 662.55 | 666.55 | 15 |  |  | 704.57 | 708.57 | 15 |  | 762.60 | 766.60 | 15 |
| 608.53 | 612.53 | 15 |  | 666.55 | 670.55 | 15 |  |  | 708.57 | 712.57 | 15 |  | 766.60 | 770.60 | 15 |
| 612.53 | 616.53 | 15 |  | 670.55 | 674.56 | 15 |  |  | 712.57 | 716.58 | 15 |  | 770.60 | 774.60 | 15 |
| 616.53 | 620.53 | 15 |  | 674.56 | 678.56 | 15 |  |  | 716.58 | 720.58 | 15 |  | 774.60 | 778.60 | 15 |
| 620.53 | 624.53 | 15 |  | 678.56 | 682.56 | 15 |  |  | 720.58 | 724.58 | 15 |  | 778.60 | 782.61 | 15 |
| 624.53 | 628.54 | 15 |  | 682.56 | 686.56 | 15 |  |  | 724.58 | 728.58 | 15 |  | 782.61 | 786.61 | 15 |
| 628.54 | 632.54 | 15 |  | 686.56 | 690.56 | 15 |  |  | 728.58 | 732.58 | 15 |  | 786.61 | 790.61 | 15 |
| 632.54 | 636.54 | 15 |  | 690.56 | 694.57 | 15 |  |  | 732.58 | 736.58 | 15 |  | 790.61 | 794.61 | 15 |
| 636.54 | 640.54 | 15 |  | 694.57 | 698.57 | 15 |  |  | 736.58 | 740.59 | 15 |  | 794.61 | 798.61 | 15 |
| 640.54 | 644.54 | 15 |  | 698.57 | 702.57 | 15 |  |  | 740.59 | 744.59 | 15 |  | 798.61 | 802.61 | 15 |
| 644.54 | 648.54 | 15 |  |  |  |  |  |  | 744.59 | 748.59 | 15 |  |  |  |  |
| 648.54 | 652.55 | 15 |  |  |  |  |  |  | 748.59 | 752.59 | 15 |  |  |  |  |
| 652.55 | 656.55 | 15 |  |  |  |  |  |  | 752.59 | 756.59 | 15 |  |  |  |  |
| 656.55 | 660.55 | 15 |  |  |  |  |  |  | 756.59 | 760.60 | 15 |  |  |  |  |
| 660.55 | 664.55 | 15 |  |  |  |  |  |  | 760.60 | 764.60 | 15 |  |  |  |  |
| 664.55 | 668.55 | 15 |  |  |  |  |  |  | 764.60 | 768.60 | 15 |  |  |  |  |
| 668.55 | 672.56 | 15 |  |  |  |  |  |  | 768.60 | 772.60 | 15 |  |  |  |  |
| 672.56 | 676.56 | 15 |  |  |  |  |  |  | 772.60 | 776.60 | 15 |  |  |  |  |
| 676.56 | 680.56 | 15 |  |  |  |  |  |  | 776.60 | 780.60 | 15 |  |  |  |  |
| 680.56 | 684.56 | 15 |  |  |  |  |  |  | 780.60 | 784.61 | 15 |  |  |  |  |
| 684.56 | 688.56 | 15 |  |  |  |  |  |  | 784.61 | 788.61 | 15 |  |  |  |  |
| 688.56 | 692.56 | 15 |  |  |  |  |  |  | 788.61 | 792.61 | 15 |  |  |  |  |
| 692.56 | 696.57 | 15 |  |  |  |  |  |  | 792.61 | 796.61 | 15 |  |  |  |  |
| 696.57 | 700.57 | 15 |  |  |  |  |  |  | 796.61 | 800.61 | 15 |  |  |  |  |
| 598.52 | 602.52 | 15 |  |  |  |  |  |  | 698.57 | 702.57 | 15 |  |  |  |  |
| 602.52 | 606.53 | 15 |  |  |  |  |  |  | 702.57 | 706.57 | 15 |  |  |  |  |
| 606.53 | 610.53 | 15 |  |  |  |  |  |  | 706.57 | 710.57 | 15 |  |  |  |  |
| 610.53 | 614.53 | 15 |  |  |  |  |  |  | 710.57 | 714.57 | 15 |  |  |  |  |
| 614.53 | 618.53 | 15 |  |  |  |  |  |  | 714.57 | 718.58 | 15 |  |  |  |  |
| 618.53 | 622.53 | 15 |  |  |  |  |  |  | 718.58 | 722.58 | 15 |  |  |  |  |
| 622.53 | 626.53 | 15 |  |  |  |  |  |  | 722.58 | 726.58 | 15 |  |  |  |  |
| 626.53 | 630.54 | 15 |  |  |  |  |  |  | 726.58 | 730.58 | 15 |  |  |  |  |
| 630.54 | 634.54 | 15 |  |  |  |  |  |  | 730.58 | 734.58 | 15 |  |  |  |  |
| 634.54 | 638.54 | 15 |  |  |  |  |  |  | 734.58 | 738.59 | 15 |  |  |  |  |
| 638.54 | 642.54 | 15 |  |  |  |  |  |  | 738.59 | 742.59 | 15 |  |  |  |  |
| 642.54 | 646.54 | 15 |  |  |  |  |  |  | 742.59 | 746.59 | 15 |  |  |  |  |
| 646.54 | 650.55 | 15 |  |  |  |  |  |  | 746.59 | 750.59 | 15 |  |  |  |  |
| 650.55 | 654.55 | 15 |  |  |  |  |  |  | 750.59 | 754.59 | 15 |  |  |  |  |

| Start | End | CES |  | Start | End | CES |  | Start | End | CES |  | Start | End | CES |
| --- | --- | --- | --- | --- | --- | --- | --- | --- | --- | --- | --- | --- | --- | --- |
| 796.61 | 800.61 | 15 |  | 854.64 | 858.64 | 15 |  | 896.66 | 900.66 | 15 |  | 954.68 | 958.69 | 15 |
| 800.61 | 804.62 | 15 |  | 858.64 | 862.64 | 15 |  | 900.66 | 904.66 | 15 |  | 958.69 | 962.69 | 15 |
| 804.62 | 808.62 | 15 |  | 862.64 | 866.64 | 15 |  | 904.66 | 908.66 | 15 |  | 962.69 | 966.69 | 15 |
| 808.62 | 812.62 | 15 |  | 866.64 | 870.65 | 15 |  | 908.66 | 912.66 | 15 |  | 966.69 | 970.69 | 15 |
| 812.62 | 816.62 | 15 |  | 870.65 | 874.65 | 15 |  | 912.66 | 916.67 | 15 |  | 970.69 | 974.69 | 15 |
| 816.62 | 820.62 | 15 |  | 874.65 | 878.65 | 15 |  | 916.67 | 920.67 | 15 |  | 974.69 | 978.69 | 15 |
| 820.62 | 824.62 | 15 |  | 878.65 | 882.65 | 15 |  | 920.67 | 924.67 | 15 |  | 978.69 | 982.70 | 15 |
| 824.62 | 828.63 | 15 |  | 882.65 | 886.65 | 15 |  | 924.67 | 928.67 | 15 |  | 982.70 | 986.70 | 15 |
| 828.63 | 832.63 | 15 |  | 886.65 | 890.65 | 15 |  | 928.67 | 932.67 | 15 |  | 986.70 | 990.70 | 15 |
| 832.63 | 836.63 | 15 |  | 890.65 | 894.66 | 15 |  | 932.67 | 936.68 | 15 |  | 990.70 | 994.70 | 15 |
| 836.63 | 840.63 | 15 |  | 894.66 | 898.66 | 15 |  | 936.68 | 940.68 | 15 |  | 994.70 | 998.70 | 15 |
| 840.63 | 844.63 | 15 |  | 898.66 | 902.66 | 15 |  | 940.68 | 944.68 | 15 |  | 998.70 | 1002.71 | 15 |
| 844.63 | 848.64 | 15 |  |  |  |  |  | 944.68 | 948.68 | 15 |  |  |  |  |
| 848.64 | 852.64 | 15 |  |  |  |  |  | 948.68 | 952.68 | 15 |  |  |  |  |
| 852.64 | 856.64 | 15 |  |  |  |  |  | 952.68 | 956.68 | 15 |  |  |  |  |
| 856.64 | 860.64 | 15 |  |  |  |  |  | 956.68 | 960.69 | 15 |  |  |  |  |
| 860.64 | 864.64 | 15 |  |  |  |  |  | 960.69 | 964.69 | 15 |  |  |  |  |
| 864.64 | 868.64 | 15 |  |  |  |  |  | 964.69 | 968.69 | 15 |  |  |  |  |
| 868.64 | 872.65 | 15 |  |  |  |  |  | 968.69 | 972.69 | 15 |  |  |  |  |
| 872.65 | 876.65 | 15 |  |  |  |  |  | 972.69 | 976.69 | 15 |  |  |  |  |
| 876.65 | 880.65 | 15 |  |  |  |  |  | 976.69 | 980.70 | 15 |  |  |  |  |
| 880.65 | 884.65 | 15 |  |  |  |  |  | 980.70 | 984.70 | 15 |  |  |  |  |
| 884.65 | 888.65 | 15 |  |  |  |  |  | 984.70 | 988.70 | 15 |  |  |  |  |
| 888.65 | 892.66 | 15 |  |  |  |  |  | 988.70 | 992.70 | 15 |  |  |  |  |
| 892.66 | 896.66 | 15 |  |  |  |  |  | 992.70 | 996.70 | 15 |  |  |  |  |
| 896.66 | 900.66 | 15 |  |  |  |  |  | 996.70 | 1000.70 | 15 |  |  |  |  |
| 798.61 | 802.61 | 15 |  |  |  |  |  | 898.66 | 902.66 | 15 |  |  |  |  |
| 802.61 | 806.62 | 15 |  |  |  |  |  | 902.66 | 906.66 | 15 |  |  |  |  |
| 806.62 | 810.62 | 15 |  |  |  |  |  | 906.66 | 910.66 | 15 |  |  |  |  |
| 810.62 | 814.62 | 15 |  |  |  |  |  | 910.66 | 914.67 | 15 |  |  |  |  |
| 814.62 | 818.62 | 15 |  |  |  |  |  | 914.67 | 918.67 | 15 |  |  |  |  |
| 818.62 | 822.62 | 15 |  |  |  |  |  | 918.67 | 922.67 | 15 |  |  |  |  |
| 822.62 | 826.63 | 15 |  |  |  |  |  | 922.67 | 926.67 | 15 |  |  |  |  |
| 826.63 | 830.63 | 15 |  |  |  |  |  | 926.67 | 930.67 | 15 |  |  |  |  |
| 830.63 | 834.63 | 15 |  |  |  |  |  | 930.67 | 934.67 | 15 |  |  |  |  |
| 834.63 | 838.63 | 15 |  |  |  |  |  | 934.67 | 938.68 | 15 |  |  |  |  |
| 838.63 | 842.63 | 15 |  |  |  |  |  | 938.68 | 942.68 | 15 |  |  |  |  |
| 842.63 | 846.63 | 15 |  |  |  |  |  | 942.68 | 946.68 | 15 |  |  |  |  |
| 846.63 | 850.64 | 15 |  |  |  |  |  | 946.68 | 950.68 | 15 |  |  |  |  |
| 850.64 | 854.64 | 15 |  |  |  |  |  | 950.68 | 954.68 | 15 |  |  |  |  |

| Start | End | CES |  | Start | End | CES |  | Start | End | CES |  | Start | End | CES |
| --- | --- | --- | --- | --- | --- | --- | --- | --- | --- | --- | --- | --- | --- | --- |
| 996.70 | 1000.70 | 15 |  | 1054.73 | 1058.73 | 15 |  | 1096.75 | 1100.75 | 15 |  | 1154.77 | 1158.78 | 15 |
| 1000.70 | 1004.71 | 15 |  | 1058.73 | 1062.73 | 15 |  | 1100.75 | 1104.75 | 15 |  | 1158.78 | 1162.78 | 15 |
| 1004.71 | 1008.71 | 15 |  | 1062.73 | 1066.73 | 15 |  | 1104.75 | 1108.75 | 15 |  | 1162.78 | 1166.78 | 15 |
| 1008.71 | 1012.71 | 15 |  | 1066.73 | 1070.74 | 15 |  | 1108.75 | 1112.76 | 15 |  | 1166.78 | 1170.78 | 15 |
| 1012.71 | 1016.71 | 15 |  | 1070.74 | 1074.74 | 15 |  | 1112.76 | 1116.76 | 15 |  | 1170.78 | 1174.78 | 15 |
| 1016.71 | 1020.71 | 15 |  | 1074.74 | 1078.74 | 15 |  | 1116.76 | 1120.76 | 15 |  | 1174.78 | 1178.79 | 15 |
| 1020.71 | 1024.72 | 15 |  | 1078.74 | 1082.74 | 15 |  | 1120.76 | 1124.76 | 15 |  | 1178.79 | 1182.79 | 15 |
| 1024.72 | 1028.72 | 15 |  | 1082.74 | 1086.74 | 15 |  | 1124.76 | 1128.76 | 15 |  | 1182.79 | 1186.79 | 15 |
| 1028.72 | 1032.72 | 15 |  | 1086.74 | 1090.75 | 15 |  | 1128.76 | 1132.76 | 15 |  | 1186.79 | 1190.79 | 15 |
| 1032.72 | 1036.72 | 15 |  | 1090.75 | 1094.75 | 15 |  | 1132.76 | 1136.77 | 15 |  | 1190.79 | 1194.79 | 15 |
| 1036.72 | 1040.72 | 15 |  | 1094.75 | 1098.75 | 15 |  | 1136.77 | 1140.77 | 15 |  | 1194.79 | 1198.79 | 15 |
| 1040.72 | 1044.72 | 15 |  | 1098.75 | 1102.75 | 15 |  | 1140.77 | 1144.77 | 15 |  | 1198.79 | 1202.80 | 15 |
| 1044.72 | 1048.73 | 15 |  |  |  |  |  | 1144.77 | 1148.77 | 15 |  |  |  |  |
| 1048.73 | 1052.73 | 15 |  |  |  |  |  | 1148.77 | 1152.77 | 15 |  |  |  |  |
| 1052.73 | 1056.73 | 15 |  |  |  |  |  | 1152.77 | 1156.78 | 15 |  |  |  |  |
| 1056.73 | 1060.73 | 15 |  |  |  |  |  | 1156.78 | 1160.78 | 15 |  |  |  |  |
| 1060.73 | 1064.73 | 15 |  |  |  |  |  | 1160.78 | 1164.78 | 15 |  |  |  |  |
| 1064.73 | 1068.74 | 15 |  |  |  |  |  | 1164.78 | 1168.78 | 15 |  |  |  |  |
| 1068.74 | 1072.74 | 15 |  |  |  |  |  | 1168.78 | 1172.78 | 15 |  |  |  |  |
| 1072.74 | 1076.74 | 15 |  |  |  |  |  | 1172.78 | 1176.78 | 15 |  |  |  |  |
| 1076.74 | 1080.74 | 15 |  |  |  |  |  | 1176.78 | 1180.79 | 15 |  |  |  |  |
| 1080.74 | 1084.74 | 15 |  |  |  |  |  | 1180.79 | 1184.79 | 15 |  |  |  |  |
| 1084.74 | 1088.74 | 15 |  |  |  |  |  | 1184.79 | 1188.79 | 15 |  |  |  |  |
| 1088.74 | 1092.75 | 15 |  |  |  |  |  | 1188.79 | 1192.79 | 15 |  |  |  |  |
| 1092.75 | 1096.75 | 15 |  |  |  |  |  | 1192.79 | 1196.79 | 15 |  |  |  |  |
| 1096.75 | 1100.75 | 15 |  |  |  |  |  | 1196.79 | 1200.80 | 15 |  |  |  |  |
| 998.70 | 1002.71 | 15 |  |  |  |  |  | 1098.75 | 1102.75 | 15 |  |  |  |  |
| 1002.71 | 1006.71 | 15 |  |  |  |  |  | 1102.75 | 1106.75 | 15 |  |  |  |  |
| 1006.71 | 1010.71 | 15 |  |  |  |  |  | 1106.75 | 1110.75 | 15 |  |  |  |  |
| 1010.71 | 1014.71 | 15 |  |  |  |  |  | 1110.75 | 1114.76 | 15 |  |  |  |  |
| 1014.71 | 1018.71 | 15 |  |  |  |  |  | 1114.76 | 1118.76 | 15 |  |  |  |  |
| 1018.71 | 1022.71 | 15 |  |  |  |  |  | 1118.76 | 1122.76 | 15 |  |  |  |  |
| 1022.71 | 1026.72 | 15 |  |  |  |  |  | 1122.76 | 1126.76 | 15 |  |  |  |  |
| 1026.72 | 1030.72 | 15 |  |  |  |  |  | 1126.76 | 1130.76 | 15 |  |  |  |  |
| 1030.72 | 1034.72 | 15 |  |  |  |  |  | 1130.76 | 1134.77 | 15 |  |  |  |  |
| 1034.72 | 1038.72 | 15 |  |  |  |  |  | 1134.77 | 1138.77 | 15 |  |  |  |  |
| 1038.72 | 1042.72 | 15 |  |  |  |  |  | 1138.77 | 1142.77 | 15 |  |  |  |  |
| 1042.72 | 1046.73 | 15 |  |  |  |  |  | 1142.77 | 1146.77 | 15 |  |  |  |  |
| 1046.73 | 1050.73 | 15 |  |  |  |  |  | 1146.77 | 1150.77 | 15 |  |  |  |  |
| 1050.73 | 1054.73 | 15 |  |  |  |  |  | 1150.77 | 1154.77 | 15 |  |  |  |  |

Supplementary Table 3. 99 Variable Window Scheme applied for SWATH data acquired on the SCIEX TripleTOF 6600+

| Start | End | CES |  | Start | End | CES |  | Start | End | CES |
| --- | --- | --- | --- | --- | --- | --- | --- | --- | --- | --- |
| 399.5 | 406.5 | 0 |  | 613.5 | 619.5 | 0 |  | 875.5 | 885.5 | 0 |
| 405.5 | 412.5 | 0 |  | 618.5 | 624.5 | 0 |  | 884.5 | 894.5 | 0 |
| 411.5 | 418.5 | 0 |  | 623.5 | 629.5 | 0 |  | 893.5 | 903.5 | 0 |
| 417.5 | 424.5 | 0 |  | 628.5 | 634.5 | 0 |  | 902.5 | 914.5 | 0 |
| 423.5 | 430.5 | 0 |  | 633.5 | 639.5 | 0 |  | 913.5 | 925.5 | 0 |
| 429.5 | 436.5 | 0 |  | 638.5 | 644.5 | 0 |  | 924.5 | 936.5 | 0 |
| 435.5 | 442.5 | 0 |  | 643.5 | 649.5 | 0 |  | 935.5 | 950.5 | 0 |
| 441.5 | 448.5 | 0 |  | 648.5 | 654.5 | 0 |  | 949.5 | 964.5 | 0 |
| 447.5 | 454.5 | 0 |  | 653.5 | 660.5 | 0 |  | 963.5 | 978.5 | 0 |
| 453.5 | 459.5 | 0 |  | 659.5 | 666.5 | 0 |  | 977.5 | 992.5 | 0 |
| 458.5 | 464.5 | 0 |  | 665.5 | 672.5 | 0 |  | 991.5 | 1011.5 | 0 |
| 463.5 | 469.5 | 0 |  | 671.5 | 678.5 | 0 |  | 1010.5 | 1030.5 | 0 |
| 468.5 | 474.5 | 0 |  | 677.5 | 684.5 | 0 |  | 1029.5 | 1054.5 | 0 |
| 473.5 | 479.5 | 0 |  | 683.5 | 690.5 | 0 |  | 1053.5 | 1078.5 | 0 |
| 478.5 | 484.5 | 0 |  | 689.5 | 696.5 | 0 |  | 1077.5 | 1117.5 | 0 |
| 483.5 | 489.5 | 0 |  | 695.5 | 702.5 | 0 |  | 1116.5 | 1156.5 | 0 |
| 488.5 | 494.5 | 0 |  | 701.5 | 708.5 | 0 |  | 1155.5 | 1200.5 | 0 |
| 493.5 | 499.5 | 0 |  | 707.5 | 714.5 | 0 |  |  |  |  |
| 498.5 | 504.5 | 0 |  | 713.5 | 720.5 | 0 |  |  |  |  |
| 503.5 | 509.5 | 0 |  | 719.5 | 726.5 | 0 |  |  |  |  |
| 508.5 | 514.5 | 0 |  | 725.5 | 732.5 | 0 |  |  |  |  |
| 513.5 | 519.5 | 0 |  | 731.5 | 738.5 | 0 |  |  |  |  |
| 518.5 | 524.5 | 0 |  | 737.5 | 744.5 | 0 |  |  |  |  |
| 523.5 | 529.5 | 0 |  | 743.5 | 750.5 | 0 |  |  |  |  |
| 528.5 | 534.5 | 0 |  | 749.5 | 756.5 | 0 |  |  |  |  |
| 533.5 | 539.5 | 0 |  | 755.5 | 763.5 | 0 |  |  |  |  |
| 538.5 | 544.5 | 0 |  | 762.5 | 770.5 | 0 |  |  |  |  |
| 543.5 | 549.5 | 0 |  | 769.5 | 777.5 | 0 |  |  |  |  |
| 548.5 | 554.5 | 0 |  | 776.5 | 784.5 | 0 |  |  |  |  |
| 553.5 | 559.5 | 0 |  | 783.5 | 791.5 | 0 |  |  |  |  |
| 558.5 | 564.5 | 0 |  | 790.5 | 798.5 | 0 |  |  |  |  |
| 563.5 | 569.5 | 0 |  | 797.5 | 805.5 | 0 |  |  |  |  |
| 568.5 | 574.5 | 0 |  | 804.5 | 812.5 | 0 |  |  |  |  |
| 573.5 | 579.5 | 0 |  | 811.5 | 819.5 | 0 |  |  |  |  |
| 578.5 | 584.5 | 0 |  | 818.5 | 826.5 | 0 |  |  |  |  |
| 583.5 | 589.5 | 0 |  | 825.5 | 834.5 | 0 |  |  |  |  |
| 588.5 | 594.5 | 0 |  | 833.5 | 842.5 | 0 |  |  |  |  |
| 593.5 | 599.5 | 0 |  | 841.5 | 850.5 | 0 |  |  |  |  |
| 598.5 | 604.5 | 0 |  | 849.5 | 858.5 | 0 |  |  |  |  |
| 603.5 | 609.5 | 0 |  | 857.5 | 867.5 | 0 |  |  |  |  |
| 608.5 | 614.5 | 0 |  | 866.5 | 876.5 | 0 |  |  |  |  |

Supplementary Table 4. 8m/z staggered window scheme for AIF data acquired on the Orbitrap QE-HFX

| start | end |  | start | end |  | start | end |  | start | end |
| --- | --- | --- | --- | --- | --- | --- | --- | --- | --- | --- |
| 400.4337 | 408.4337 |  | 728.5829 | 736.5829 |  | 452.4574 | 460.4574 |  | 780.6065 | 788.6065 |
| 408.4374 | 416.4374 |  | 736.5865 | 744.5865 |  | 460.461 | 468.461 |  | 788.6102 | 796.6102 |
| 416.441 | 424.441 |  | 744.5901 | 752.5901 |  | 468.4647 | 476.4647 |  | 796.6138 | 804.6138 |
| 424.4446 | 432.4446 |  | 752.5938 | 760.5938 |  | 476.4683 | 484.4683 |  | 804.6174 | 812.6174 |
| 432.4483 | 440.4483 |  | 760.5974 | 768.5974 |  | 484.4719 | 492.4719 |  | 812.6211 | 820.6211 |
| 440.4519 | 448.4519 |  | 768.6011 | 776.6011 |  | 492.4756 | 500.4756 |  | 820.6248 | 828.6248 |
| 448.4556 | 456.4556 |  | 776.6047 | 784.6047 |  | 500.4792 | 508.4792 |  | 828.6284 | 836.6284 |
| 456.4592 | 464.4592 |  | 784.6083 | 792.6083 |  | 508.4828 | 516.4828 |  | 836.632 | 844.632 |
| 464.4628 | 472.4628 |  | 792.6119 | 800.6119 |  | 516.4865 | 524.4865 |  | 844.6356 | 852.6356 |
| 472.4665 | 480.4665 |  | 800.6156 | 808.6156 |  | 524.4901 | 532.4901 |  | 852.6393 | 860.6393 |
| 480.4701 | 488.4701 |  | 808.6193 | 816.6193 |  | 532.4938 | 540.4938 |  | 860.6429 | 868.6429 |
| 488.4738 | 496.4738 |  | 816.6229 | 824.6229 |  | 540.4974 | 548.4974 |  | 868.6465 | 876.6465 |
| 496.4774 | 504.4774 |  | 824.6265 | 832.6265 |  | 548.501 | 556.501 |  | 876.6501 | 884.6501 |
| 504.481 | 512.481 |  | 832.6302 | 840.6302 |  | 556.5046 | 564.5046 |  | 884.6538 | 892.6538 |
| 512.4846 | 520.4846 |  | 840.6338 | 848.6338 |  | 564.5083 | 572.5083 |  | 892.6575 | 900.6575 |
| 520.4883 | 528.4883 |  | 848.6375 | 856.6375 |  | 572.512 | 580.512 |  | 900.6611 | 908.6611 |
| 528.4919 | 536.4919 |  | 856.6411 | 864.6411 |  | 580.5156 | 588.5156 |  | 908.6647 | 916.6647 |
| 536.4956 | 544.4956 |  | 864.6447 | 872.6447 |  | 588.5192 | 596.5192 |  | 916.6683 | 924.6683 |
| 544.4992 | 552.4992 |  | 872.6484 | 880.6484 |  | 596.5228 | 604.5228 |  | 924.672 | 932.672 |
| 552.5028 | 560.5028 |  | 880.652 | 888.652 |  | 604.5265 | 612.5265 |  | 932.6757 | 940.6757 |
| 560.5065 | 568.5065 |  | 888.6556 | 896.6556 |  | 612.5302 | 620.5302 |  | 940.6793 | 948.6793 |
| 568.5101 | 576.5101 |  | 896.6593 | 904.6593 |  | 620.5338 | 628.5338 |  | 948.6829 | 956.6829 |
| 576.5138 | 584.5138 |  | 904.6629 | 912.6629 |  | 628.5374 | 636.5374 |  | 956.6865 | 964.6865 |
| 584.5174 | 592.5174 |  | 912.6666 | 920.6666 |  | 636.541 | 644.541 |  | 964.6902 | 972.6902 |
| 592.521 | 600.521 |  | 920.6702 | 928.6702 |  | 644.5447 | 652.5447 |  | 972.6938 | 980.6938 |
| 600.5247 | 608.5247 |  | 928.6738 | 936.6738 |  | 652.5483 | 660.5483 |  | 980.6975 | 988.6975 |
| 608.5283 | 616.5283 |  | 936.6775 | 944.6775 |  | 660.5519 | 668.5519 |  | 988.7011 | 996.7011 |
| 616.532 | 624.532 |  | 944.6811 | 952.6811 |  | 668.5556 | 676.5556 |  | 996.7047 | 1004.7047 |
| 624.5356 | 632.5356 |  | 952.6848 | 960.6848 |  | 676.5592 | 684.5592 |  |  |  |
| 632.5392 | 640.5392 |  | 960.6884 | 968.6884 |  | 684.5629 | 692.5629 |  |  |  |
| 640.5428 | 648.5428 |  | 968.692 | 976.692 |  | 692.5665 | 700.5665 |  |  |  |
| 648.5465 | 656.5465 |  | 976.6957 | 984.6957 |  | 700.5701 | 708.5701 |  |  |  |
| 656.5502 | 664.5502 |  | 984.6993 | 992.6993 |  | 708.5738 | 716.5738 |  |  |  |
| 664.5538 | 672.5538 |  | 992.7029 | 1000.7029 |  | 716.5774 | 724.5774 |  |  |  |
| 672.5574 | 680.5574 |  | 396.4319 | 404.4319 |  | 724.5811 | 732.5811 |  |  |  |
| 680.561 | 688.561 |  | 404.4355 | 412.4355 |  | 732.5847 | 740.5847 |  |  |  |
| 688.5647 | 696.5647 |  | 412.4392 | 420.4392 |  | 740.5883 | 748.5883 |  |  |  |
| 696.5684 | 704.5684 |  | 420.4428 | 428.4428 |  | 748.592 | 756.592 |  |  |  |
| 704.572 | 712.572 |  | 428.4465 | 436.4465 |  | 756.5956 | 764.5956 |  |  |  |
| 712.5756 | 720.5756 |  | 436.4501 | 444.4501 |  | 764.5992 | 772.5992 |  |  |  |
| 720.5792 | 728.5792 |  | 444.4537 | 452.4537 |  | 772.6029 | 780.6029 |  |  |  |

Supplementary Table 5. Collision energy ramp of each Gas-phase fractionated SONAR run.

| m/z start | m/z end | CE start | CE end |
| --- | --- | --- | --- |
| 400 | 500 | 16 | 20 |
| 500 | 600 | 20 | 24 |
| 600 | 700 | 24 | 28 |
| 700 | 800 | 28 | 32 |
| 800 | 900 | 32 | 36 |
| 900 | 1000 | 36 | 40 |
| 1000 | 1100 | 40 | 44 |
| 1100 | 1200 | 44 | 48 |

Supplementary Table 6. diaPASEF acquisition scheme

| Frame | Window 1 | Window 2 |
| --- | --- | --- |
| 1 | 400-426 | 800-826 |
| 2 | 425-451 | 825-851 |
| 3 | 450-476 | 850-876 |
| 4 | 475-501 | 875-901 |
| 5 | 500-526 | 900-926 |
| 6 | 525-551 | 925-951 |
| 7 | 550-576 | 950-976 |
| 8 | 575-601 | 975-1001 |
| 9 | 600-626 | 1000-1026 |
| 10 | 625-651 | 1025-1051 |
| 11 | 650-676 | 1050-1076 |
| 12 | 675-701 | 1075-1101 |
| 13 | 700-726 | 1100-1126 |
| 14 | 725-751 | 1125-1151 |
| 15 | 750-776 | 1150-1176 |
| 16 | 775-801 | 1175-1201 |

Supplementary Table 7. Overview of the amount of E.coli protein digest loaded for each AutoQC acquisition.

| Instrument | Sample Loading |
| --- | --- |
| Sciex TripleTOF 5600 (capillary flow) | 800 ng |
| Sciex TripleTOF 6600+ (capillary flow) | 800 ng |
| Thermo Orbitrap QE-HFX (nano flow) | 400 ng |
| Waters Synapt G2-Si (nano flow) | 400 ng |
| Waters Synapt XS (capillary flow) | 800 ng |
| Bruker TimsTOF Pro (nano flow) | 400 ng |
